# Opposing patterns of abnormal D1 and D2 receptor dependent cortico-striatal plasticity explain increased risk taking in patients with DYT1 dystonia

**DOI:** 10.1101/869743

**Authors:** Tom Gilbertson, David Arkadir, J. Douglas Steele

**Author notes:** Corresponding author: Dr. Tom Gilbertson, Department of Neurology, Ninewells Hospital & Medical School, Level 6, South Block, Dundee, DD1 9SY,.

## Abstract

Patients with dystonia caused by the mutated TOR1A gene exhibit a risk neutral behaviour compared to controls who are risk averse in the same reinforcement learning task. We hypothesised this increased risk taking could be reproduced by a reinforcement learning model which included biologically realistic striatal plasticity learning rules. We aimed to test whether a specific combination of cortico-striatal plasticity abnormalities at D1 and D2 receptors could explain the abnormal behaviour. We found a model of cortico-striatal plasticity could generate simulated behaviour indistinguishable from patients only when both D1 and D2 plasticity was abnormally increased in opposite directions: specifically when D1 synaptic potentiation and D2 depotentiation were both increased. This result is consistent with previous observations in rodent models of cortico-striatal plasticity at D1 receptors, but contrasts with the pattern reported *in vitro* for D2 synapses. This suggests that additional factors in patients who manifest motor symptoms may lead to divergent effects on D2 synaptic plasticity that are not apparent in rodent models of this disease.

## Introduction

Cortico-striatal plasticity has been implicated in the acquisition and extinction of learned actions through positive (Reynolds *et al.*, 2001) and negative reinforcement learning (Dalley *et al.*, 2007). Optogenetic studies have confirmed a causal role for phasic dopamine in the form of the reward prediction error signal in determining behavioural choices (Kravitz *et al.*, 2012; Steinberg *et al.*, 2013; Chang *et al.*, 2016). This has lead to the widely accepted view that dopamine modifies behaviour by mediating its opposing effects on cortico-striatal synaptic strength via the two subtypes of dopamine receptor (Shen *et al.*, 2008). Within this framework, the positive-going prediction error signal, which accompanies a rewarding outcome, strengthens cortico-striatal synapse within the “direct” or striato-nigral pathway and increases the likelihood of this choice being repeated. Conversely, the negative-going prediction error signal associated with an aversive outcome leads to strengthening of the “indirect” or striato-pallidal pathway, which suppresses the likelihood of choice repetition. Both of these signals rely upon the induction of cortico-striatal long-term potentiation (LTP) to mediate their behavioural effect, albeit under opposite dopaminergic conditions and via D1 and D2 receptors(Frank, 2005). Accordingly, in humans, individual sensitivity to positive and negative feedback (via positive and negative going prediction error signals) correlates with the extent of D1 or D2 receptor (D1R and D2R respectively) expression and genetic influences on their variability (Cools *et al.*, 2009; Frank *et al.*, 2009; Cox *et al.*, 2015)

The mutated *TOR1A* gene causes generalised dystonia (DYT1), a movement disorder characterised by sustained or intermittent muscle contractions leading to abnormal repetitive movements and postures (Ozelius *et al.*, 1997). Brain slice recordings from rodents expressing the human mutant gene exhibit abnormal cortico-striatal plasticity with a combination of abnormally strong LTP; (Martella *et al.*, 2009) and weak long-term depression, LTD; (Martella *et al.*, 2009; Grundmann *et al.*, 2012). Subsequent studies have delineated a receptor specific post-synaptic abnormality in D2R transmission as the principle cause for impaired LTD at the cortico-striatal synapse (Napolitano *et al.*, 2010). In view of the importance of cortico-striatal plasticity in reinforcement learning, Arkadir et. al., proposed that patients with the TOR1A mutation should exhibit a learning strategy that is contingent with the abnormal plasticity seen in rodent models (Arkadir *et al.*, 2016). The patients in this study were found to be significantly more likely to make a risky choice in a reinforcement learning task compared to controls. They concluded that this risk taking behaviour was consistent with asymmetric integration of the positive and negative prediction error signals as a consequence of maladaptive striatal plasticity. Given the distinct effects that these signals mediate on the direct and indirect pathways, they proposed three candidate D1R and D2R abnormalities that may lead to the pattern of behaviour observed: 1) An increased sensitivity to a “win,” due to abnormally increased D1R mediated LTP with intact D2R signalling, 2) increased sensitivity to a “win,” with blunted sensitivity to a “loss” both due to abnormally increased D1R - LTP and D2R-LTD, 3) abnormally increased D1R-LTP and D2-LTP with blunted LTD at both receptors. The third explanation was favoured as it was consistent with the pattern observed from the rodent slice data. This pattern is nevertheless the most difficult of the three to reconcile with increased risk taking behaviour. If this were indeed the underlying cause, any increased riskiness mediated by pathological D1-LTP would be working in opposition to the risk aversive effects of increased D2-LTP. In this scenario, increased risk taking could therefore only be conferred by a D1 abnormality that was substantially greater than the D2 abnormality.

We wanted to address this conflict between the reported plasticity abnormalities demonstrated in rodent models and the risk taking behaviour observed in patients using a model of cortico-striatal plasticity (Gilbertson *et al.*, 2019). In these simulations the model reproduced decision making in the task whilst being forced to learn under the three proposed conditions of striatal plasticity. We found the pattern of D1R/D2R abnormality from the rodent experiments was least robust at reproducing actual experimental behaviour of patients. In contrast, the model generated simulated behaviour that was statistically indistinguishable from that observed experimentally by patients, only when learning under conditions where the opposite pattern of D2R abnormality (reduced LTP / increased LTD) was present. We propose this abnormality is easily reconciled with current understanding of the neurobiology of learning and increased risk taking. Notably, we suggest that D2R dysfunction may fundamentally differ between dystonically manifest patients and non-dystonically manifest animal models which share the *TOR1A* gene mutation.

## Methods

### Subjects and behavioural paradigm

Behavioural data was from Arkadir et. al., (2016) which included 13 adult patients with DYT1 dystonia and 13 age and sex-matched controls. Further details regarding their medications and clinical assessments are described in detail in the original manuscript. The trial-and-error (reinforcement) learning task consisted of 326 trial presentations of four pseudo-letters which served as cues (‘slot machines’). This included an initial familiarisation (training phase) of 26 trials. Each cue was attributed a different reward schedule (sure 0¢, sure 5¢, sure 10¢, and the so-called “‘risky” cue associated with 50:50% probabilities of 0¢ or 10¢ payoffs). The task consisted of pseudo-randomised presentations of the cues in either “forced” or “choice” trials (Figure 1). Pay-out feedback was presented either following a “forced,” trial when one of the four cues was presented on its own and selected. During a “choice” trial, feedback was given following the subject’s choice of one cue from a pair presented. One of five pairs of cue combinations were presented during the “choice” trials. These included 0¢ versus 5¢, 5¢ versus 10¢, 0¢ versus 0/10¢, 5¢ versus 0/10¢ and 10¢ versus 0/10¢. The principle behavioural result reported by Arkadir et. al., was an increased tendency for patients to choose the risky cue when presented with the 5¢ versus 0/10¢ pairing. We therefore focused our re-analysis of their data on these “risk” choice trials highlighted by Arkadir et. al. To ensure consistency with their analysis of the task behaviour, we report in an identical fashion, the overall proportion of risky cue choice both across the task (n = 60 trials) as a whole (Figure 1A) and across four (n=15 trial) blocks (Figure 1B).

**Figure 1:**
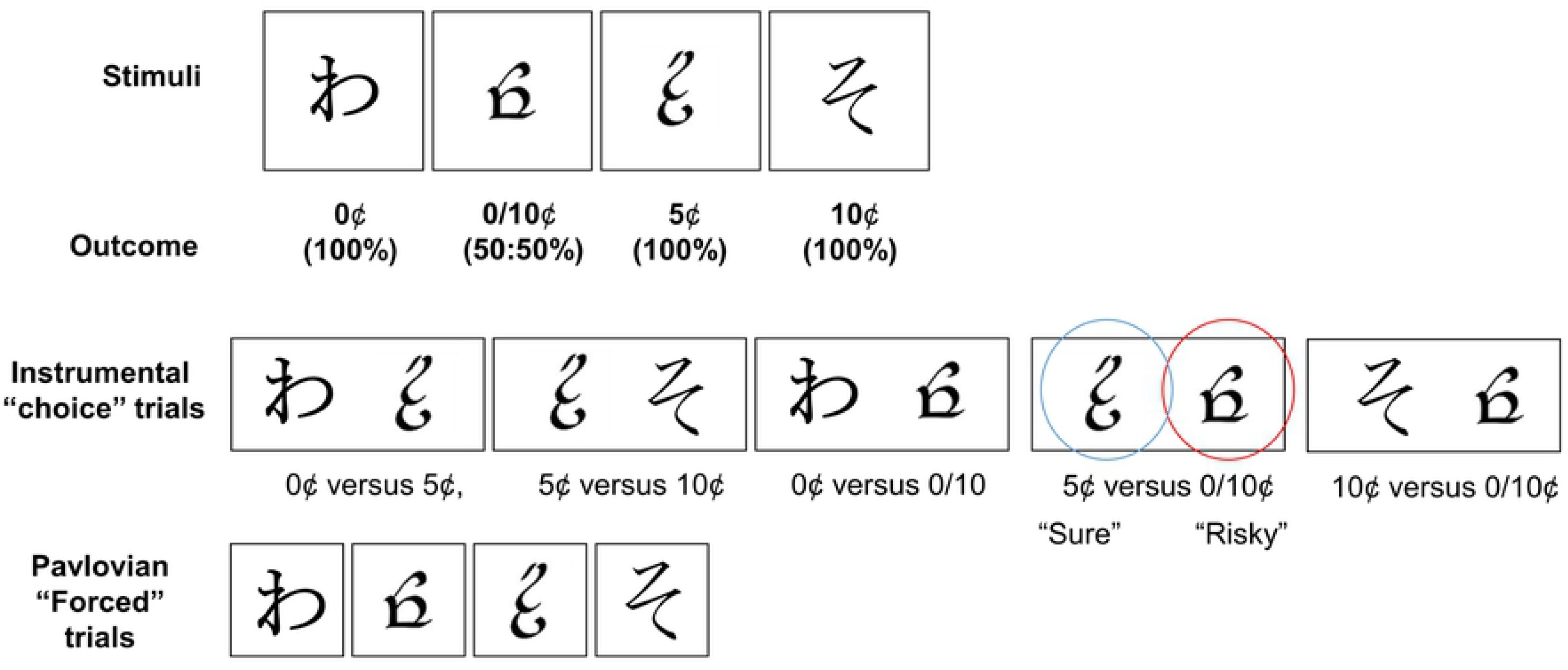
Task. Examples of visual stimuli used in the reinforcement learning task by Arkadir et. al., 2016. Trials were randomly presented as either single stimuli which required a forced choice and corresponding outcome or as instrumental trials where subjects were instructed to choose one of two of the stimuli. The risky cue choice trials were between the “risky cue” whose choice led to a 50% chance of 10¢ or 0¢ (highlighed here by the red circle) or the “sure cue” which had a 100% chance of 5¢ payout.

### Model fitting

The behavioural data was fitted to the cortico-striatal plasticity (CSP) model described in detail in Gilbertson et. al. (2019). This combines both traditional temporal difference (TD) models of reinforcement learning with biologically plausible cortico-striatal synaptic weight changes based upon *in vitro* data (Shen *et al.*, 2008). At the core of this model are two striatal populations, representing the D1R and D2R expressing direct and indirect pathways. The output of these are in turn a function of the interaction between the reward prediction error (RPE) signal (*R*(*t*) − *Q*(*A*, *t* − 1)) in the equation;

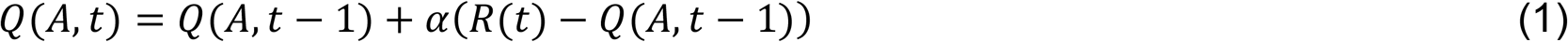

Where *α* is the learning rate, R_t_ is the outcome (reward[1] or nothing[0]), and the striatal activity *S*_*n*_of each population on trial *t* for action *A* was defined as:

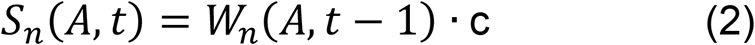

where *W* is the cortico-striatal synaptic weight and c is a constant input of 1. Here we assume two striatal “populations” *S*_*D*1_ and *S*_*D*2_; the D1 receptor expressing direct and D2 receptor expressing indirect pathways respectively. Each population represents four actions (corresponding to the four cue choices in the task). The cortico-striatal synaptic weights in each population are modified at the synapse corresponding to the chosen action *A;*

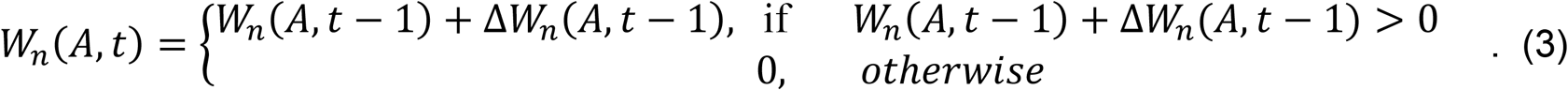

With the change in synaptic weight being the product of the striatal postsynaptic activity and the influence of dopamine:

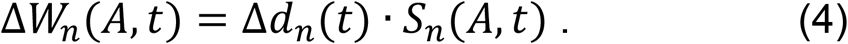

Here the magnitude Δ*d*_*n*_ of dopamine’s effect on synaptic plasticity is,

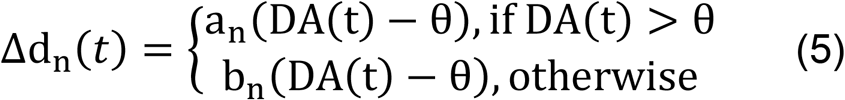

where (*a*_*n*_,*b*_*n*_) are coefficients determining the dependence of synaptic plasticity on the current trial’s level of dopamine DA(t), and the constant *θ* determines the baseline level of dopamine. Equation 5 links the RPE from Equation 1 by;

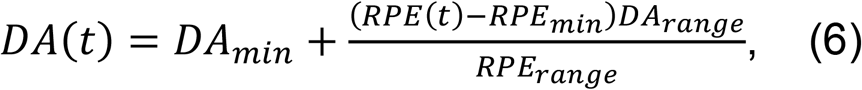

where RPE(t) < 0, *DA*_*min*_ = 0, *DA*_*range*_ = *θ*, *RPE*_*min*_ = −1, *RPE*_*range*_ = 1; otherwise *DA*_*min*_ = *θ*, *DA*_*range*_ = 1 − *θ*, *RPE*_*min*_ = 0, *RPE*_*range*_ = 1.

For forced trials the striatal population’s weight *W*_*n*_(*A*, *t*) is updated for the forced action choice only. During choice trials the models chosen action is determined by competition between the two striatal pathways for control of the pallidal output. The striatal weights are then updated for the action chosen from the pair of choices. Thus, for a choice trial with two actions(*A*_1_, A_2_);

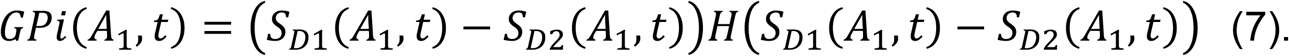

where *H()* is the Heaviside step function: *H(x)* = 0 if *x* ≤ 0, and *H(x)* = 1 otherwise; and similarly for action *A*_2_. In turn the probability of choosing action A_1_ was determined by the softmax equation with the basal ganglia’s output substituted for the value term:

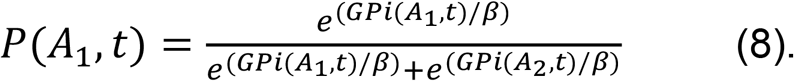

The CSP model requires estimation of six free parameters. This includes two relating to the phasic dopamine (RPE) signal, namely the learning rate (*α*) and reward sensitivity or inverse temperature parameter (*β*), and four parameters which govern the magnitude of D1 and D2 mediated CS plasticity; *a*_*1*_ (D1-LTD) *b*_*1*_ (D1-LTP), *a*_*2*_ (D2-LTP), *b_2_* (D2-LTD). Each of these parameters govern the gradient of the synaptic weight change function and its interaction with phasic dopamine. Larger values of each parameter lead to more significant changes in synaptic weight across the dynamic range of dopamine, as this is encoded in the positive and negative prediction error signals.

Estimation of the 6 parameters (*a*_*1*_, *b*_*1*_, *a*_*2*_, *b*_*2*_, *α*, *β*) was performed simultaneously using data from the whole task including all trials of both types (forced and choice) and the initial training phase. We optimised the model parameters by minimising the negative log likelihood of the data given each parameter combination. This was done using the Matlab (Mathworks, NA) function *fmincon.* The initial starting points of this function were estimated following a grid-search of the parameter space. The bounds of both *fmincon* and the grid-search were defined as *a*_*1*_ = [0,2.5]; *b*_*1*_ = [0, 1.5]; *a*_*2*_ = [−2.5,0], *b*_*2*_ = [−1.5,0], *α* = [0, 1], *β* = [0,2]. (The softmax equation in the CSP model divides by beta hence the range here has low values relative to TD models where beta multiplies). The intervals for the grid-search were 0.2, for the “*a*” parameters, 0.1 for the “*b*” parameters and 0.1 and 0.2 for *α* and *β* respectively due to allow for the differences in the ranges of their bounds.

Probability density functions for each of the four plasticity parameters were generated by fitting a nonparametric kernel function to control subject’s estimates. These were used to determine the parameter space bounds that defined “pathologically” high (>95%) or low (<5%) plasticity within the model’s parameter space. For hypotheses testing where the plasticity was considered to be within the normal “physiological” range, the bounds were defined by the 5% and 95% confidence limits of the control subject values. Fitting was then performed separately for each hypothesis (**H1**-**H3**) in turn. For simplicity we label these as:- “**H1**” Increased D1-LTP & decreased D1-LTD, “**H2**” Increased D1-LTP & decreased D1-LTD, Decreased D2-LTP & increased D2-LTD, “**H3**” Increased D1-LTP & decreased D1-LTD, Increased D2-LTP & decreased D2-LTD.

## Results

### Controls

To test the reliability of the final model fitting and its ability to capture healthy control behaviour, experimental data sets (n=1000) were simulated, using the final parameter estimates (See Table 1 for values). These simulations were generated using the final individual subject parameters incorporated into the CSP model re-performing the task with the original experimental cue sequence. We compared the simulated model decisions to choose the risky cue to the choice probabilities from the control subject’s experimental data, by performing a two-way ANOVA with two independent variables: source of choices [e.g. simulation, experiment], block number [1-4]. There was no significant difference in the probability of choosing the risky cue in the experimental behavioural data or the simulated behavioural data (ANOVA, F (1) = 0.01, p=0.91), or any difference between the simulated or experimental risky choices across the four blocks of the tasks (ANOVA [Source, Block], F (3) = 0.4. P = 0.75). For an illustrative comparison, the experimental probabilities of choosing the risky cue are plotted in blue for the controls in Figure 2, with both experimental and simulated choices overlaid in Figure 3. This analysis suggests that the average choice behaviour between each block in the task could be simulated using the CSP model for individual controls, and that this was statistically indistinguishable from that seen experimentally.

**Figure 2:**
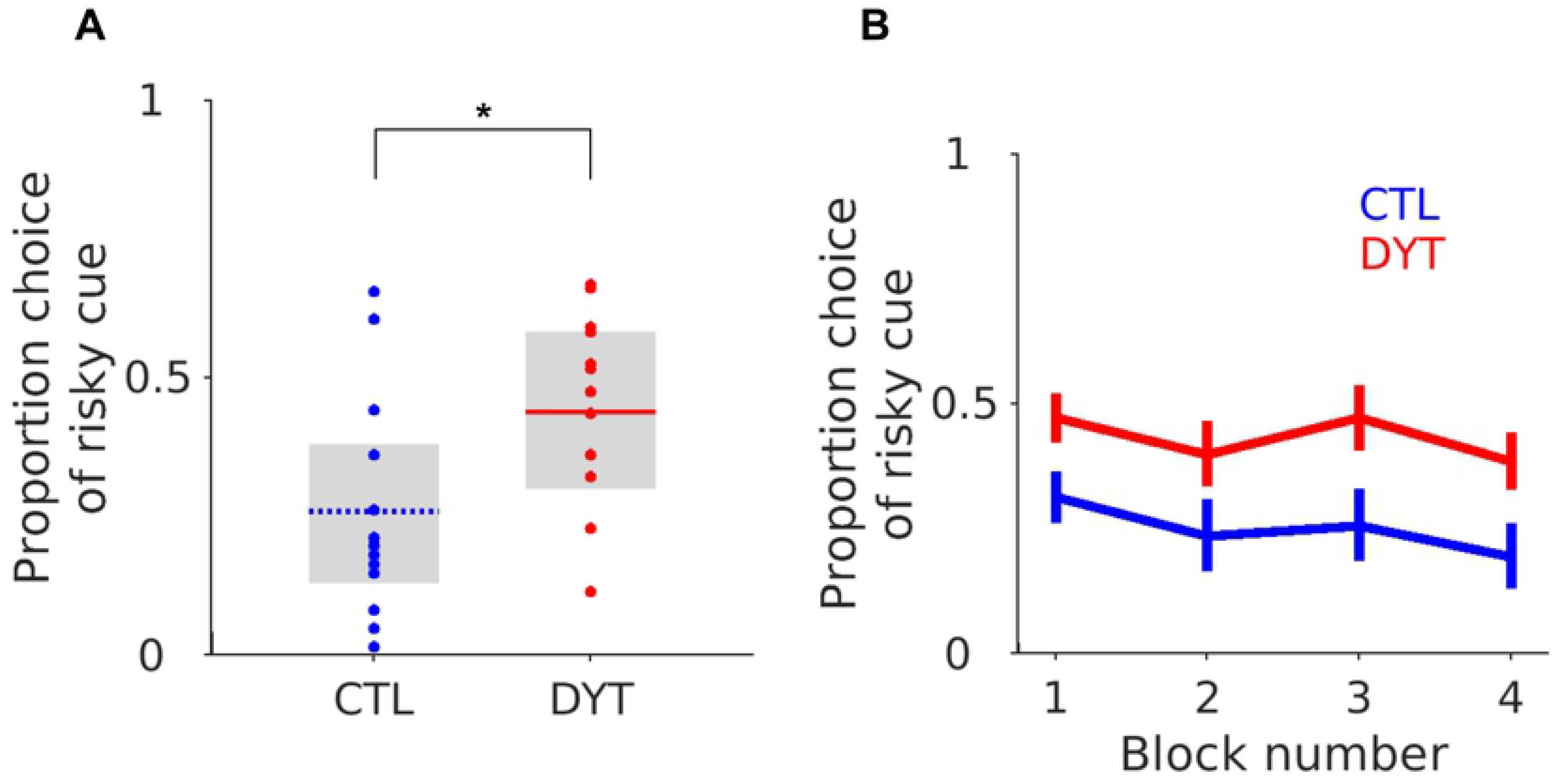
Experimental risk taking behaviour in patients and controls. (**A**) Boxplots illustrate the mean choice probability of the patients (red) and controls (blue) represented by the horizontal lines across the task as a whole. Each individual subjects choice probabilities are superimposed. The grey boxes represent the interquartile range. (**B**) The patients and controls average choice probabilities across four 15 trial blocks over the course of the task. The error bars represent the S.E.M. * Mann-Whitney z = 2.33, P < 0.05.

**Figure 3:**
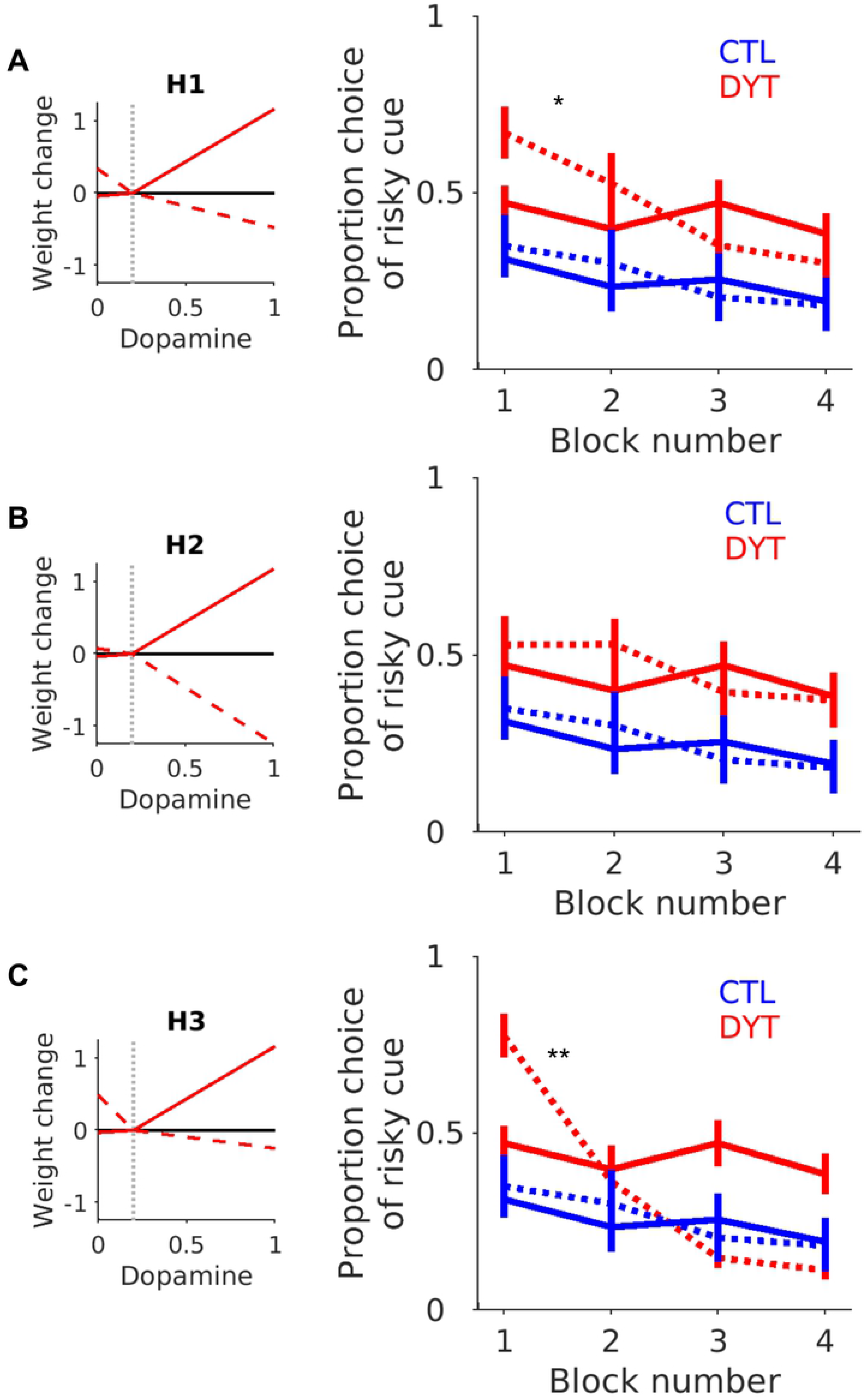
Simulated risk taking under each hypothetical plasticity abnormality. Each plot from **A**-**C**illustrates the final average synaptic weight change curve for the patients under each hypothetical plasticity condition (H1-H3). See text for details of the D1 and D2 receptor abnormality for each hypothesis. The average simulated (n=1000 simulations) choice probability of the risky cue for each block (1-4) in the task is represented by the dashed (--) lines with the patients in red and controls in blue. The error bars represent the average standard error across the simulated experiments. The solid lines (-) represent the average choice (±S.E.M) from the experimental data of Arkadir et. al. (2016). Significant differences between the simulated and experimental mean choice probability were present under plasticity conditions for H1 (*p=0.01) and H3 (**p<0.001) but not for H2, consistent with the overlapping experimental and simulated choices for this hypothesis.

### Patients

Re-analysing the experimental data of Arkadir et al., we found the same tendency for patients to show significantly less risk aversion (Figure 2A), choosing the risky stimulus significantly more often than controls (DYT 0.44 ± 0.04, CTL 0.26 ± 0.05, Mann-Whitney z = 2.23, df = 24, P < 0.05). Importantly, the patients increased risky decision taking continued throughout the four experimental blocks (conducting a one-way ANOVA with task block as a single independent variable, demonstrated no significant effect of block, F (1) =0.62, p=0.61). Recalling that the choice of the risky cue led to a 50:50% probability of either 0 or 10$ outcome, this absence of any modification of risk taking behaviour over time is despite receiving proportionately more 0$ (losing) outcomes (Figure 2B). Our aim of fitting the patient’s behaviour data was therefore to capture both the overall level of riskiness across the task and this absence of risky cue devaluation between blocks. We therefore re-fitted the patient’s behavioural data whilst constraining the bounds of the fitting procedure to the parameter space defined by the three hypothesised plasticity combinations (H1-H3: “**H1**” Increased D1-LTP & decreased D1-LTD, “**H2**” Increased D1-LTP & decreased D1-LTD, Decreased D2-LTP & increased D2-LTD, “**H3**” Increased D1-LTP & decreased D1-LTD, Increased D2-LTP & decreased D2-LTD). Comparing the individual negative log likelihoods of each hypothesis demonstrated a trend towards H1 and H2 (10 subjects) explaining the behaviour better than H3 (Fisher exact test x^2^ (24)=11.1, p =0.05 Bonferroni corrected), but no overall single wining hypothesis. Given the similarity of both the negative log likelihood values and the overlap between the hypothetical plasticity abnormalities, we tested whether any one of the hypothesis could recover the risky choice behaviour by comparing their simulated (generated) risk taking behaviour. We generated simulated “experiments” (n=1000) using the individual patients parameter estimates for each hypothesis fitted. The results are plotted alongside the simulated and experimental control data in Figure 3. As illustrated (Figure 3B) the only hypothesis, which could accurately recover the experimental behaviour, was H2 (Increased D1-LTP & decreased D1-LTD, Decreased D2-LTP & increased D2-LTD). A feature of the alternative hypotheses (H1 & H3) was their inability to capture the between-block risk taking behaviour of the patients which remained relatively similar across the whole task (i.e. from blocks 1-4 the risky cue was chosen to a similar degree). In contrast, when the model performed the task with the predefined plasticity abnormalities associated with H1 & H3, the models choice probability of the risky cue substantially reduced between the beginning (block 1) and end of the task (block 4).

A feature of the experimental patient’s behaviour was a combination of both a raised baseline level of risky choice and an absence of any devaluation of the risky cue between blocks as the experiment progressed. Only under the conditions of H2’s plasticity combination (increased D1 “go” & decreased D2-NoGo) could this more risky and non-devaluing between-block combination of behaviours be reproduced by the model simulations. Statistically, this observation was reflected by there being no discernible difference between the simulated model and the experimental patient’s risky cue choice probability. Conducting a two-way ANOVA with two independent variables (source of choices [simulation or experiment], task block), there was no effect of the source of the choice data (ANOVA, F (1) = 0.44, p=0.50) or any significant interaction between the variables (ANOVA, F(3) = 1.48, p =0.21). Consistent with the experimental choice behaviour in the patients, there was no statistically significant between-block differences in choice probability for the simulations under H2’s plasticity conditions (ANOVA, F(3)=1.99, p=0.12). In contrast, there was a significant difference in the simulated decision making of the model under the plasticity conditions of H1 and H3. For both hypotheses there was a significant interaction between the variables for H1 (ANOVA, F (3) = 3.63, p = 0.01) and H3, (ANOVA, F (3) = 32.12, p<0.001). Furthermore, there was also an effect of block for both hypotheses, H1, (ANOVA, F(3) = 5.46, p<0.01), H3 (ANOVA, F(3) = 43.49, p<0.001). The choice probability across the task for both of these models therefore contrasted with and did not capture, the experimental patient behaviour where no statistical difference was detected between each block of the task (see above). In all, this analysis would support the assertion that the only hypothesis that could accurately reproduce both the risk neutral behaviour of the patients and their behaviour between blocks across the task, was the combined increased D1-LTP to LTD and decreased D2-LTP to LTD. The reliability of the model under the plasticity conditions of H2 to replicate the experimental behaviour is further illustrated in Figure 4A. Here we plot a single simulated experiment and for illustrative proposes, a random sample of 100 (from the 1000 generated) simulated control and dystonia behavioural experiments.

**Figure 4:**
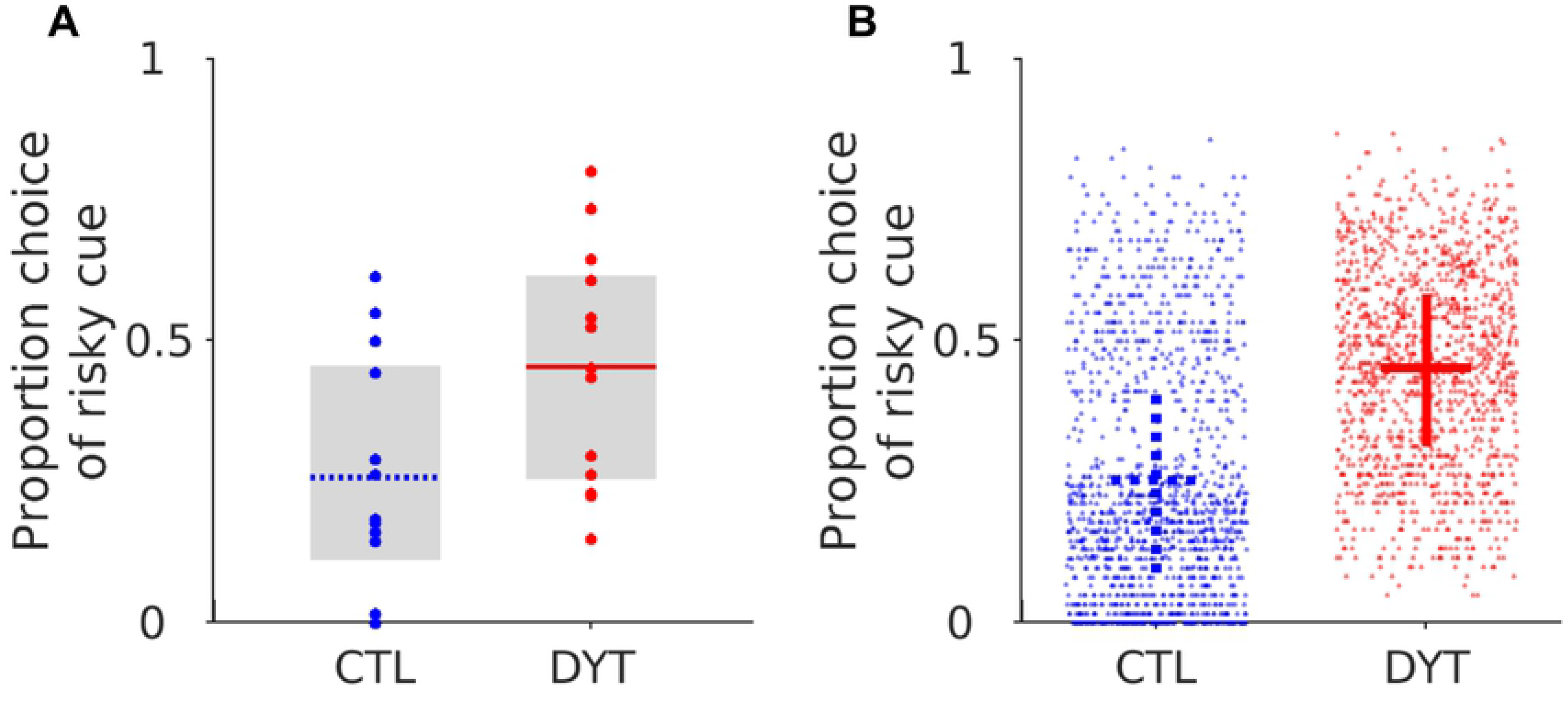
Simulated choices under plasticity conditions for H2. Example of a single simulated experiment using the final parameters estimates for the controls and patient estimates with H2 (**A**). This captures both the experimental mean and individual variance in both groups and closely replicates the experimental behaviour The CSP model was robust in replicating this behaviour across multiple simulations (**B**). For illustrative purposes we plot the first 100 of the 1000 simulated data sets from both the individual controls (blue) and patients (red). The mean choice of the risky cue and interquartile range (average between simulation) are represented by the dashed blue and solid red cross-hairs in the controls and patients respectively.

Consistent with the constraints on the fitting procedure for H2, where all four parameters were in the “pathological” range, the final plasticity parameters fitted to the patients (*a*_*1*_-*b*_*2*_) were all significantly different to the healthy controls (Two-way ANOVA (F(3) = 30, p<0.001). In contrast, there was no corresponding difference in the *α* (Mann-Whitney z (24) = 1.2, p=0.23), or *β* terms (Mann-Whitney z (24) = 1.2, p=0.22). The final dopamine weight change curve for patients (H2) and controls illustrates the expected effects of dopamine in the presence of increased D1-LTP to LTD and decreased D2-LTP to LTD (Figure 5). Relative to controls, patients significantly strengthened the direct pathway (D1R expressing) and weakened the cortico-striatal synaptic connection in the indirect pathway,(D2R expressing), in response to the positive prediction error (dopamine burst). Conversely, the negative prediction error following a loss (and corresponding dip in dopamine) produced less D2-LTP and D1-LTD and overall less risk aversive choice behaviour.

**Figure 5:**
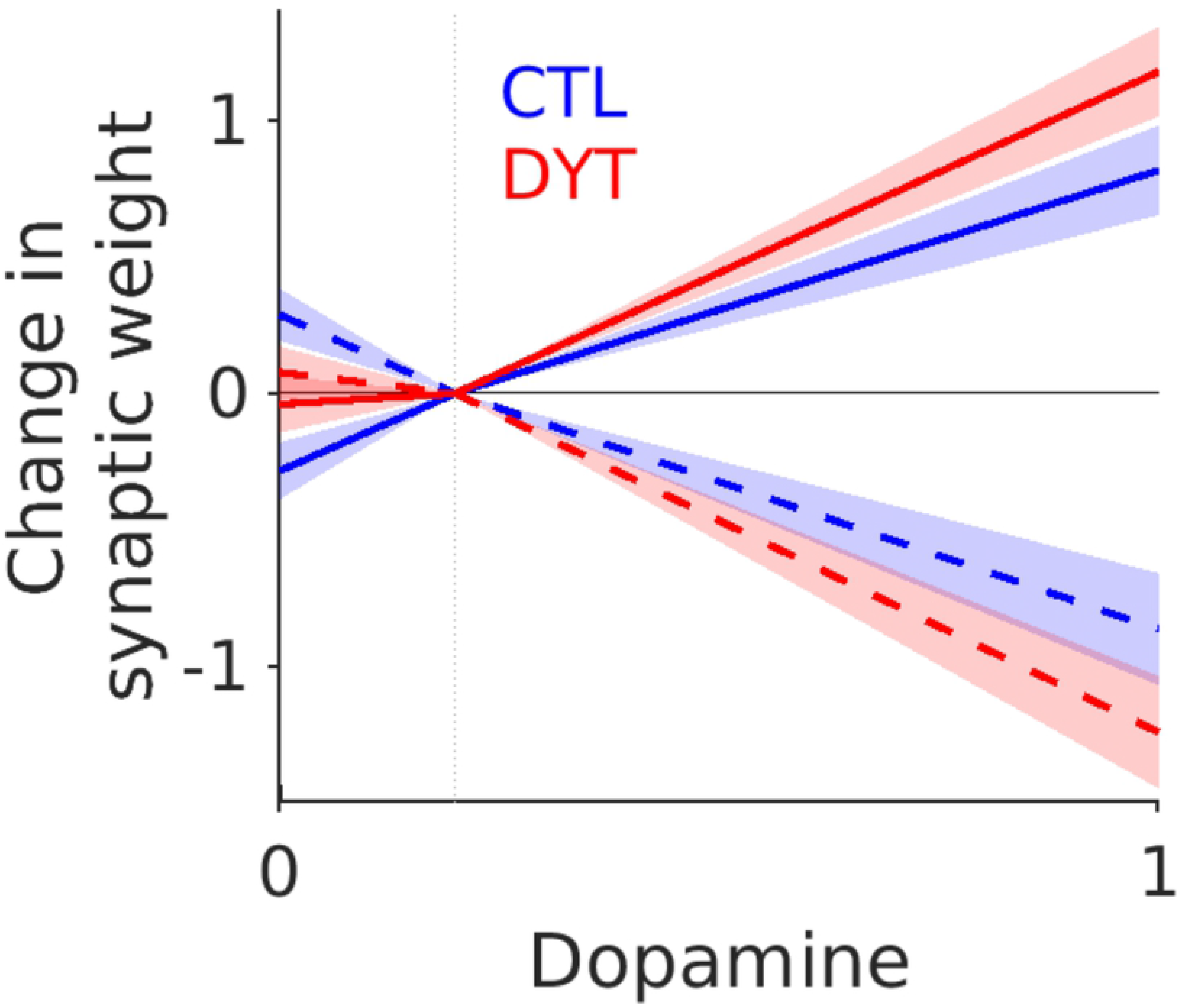
Final dopamine-synaptic weight change for patients and controls. Solid lines, D1, dashed lines D2. Mean values ± S.E.M represented by shaded area. Patients in red, controls in blue.

To understand why the CSP model could only recover the behaviour of the patients when both plasticity in the direct (D1) and indirect (D2) pathways were affected in opposite directions, we examined the time course of changes in D1 and D2 synaptic plasticity in the model through the task. These are illustrated for H1 & H2 in Figure 5A and B. As expected for a striatum where D1 plasticity is biased towards synaptic potentiation, the synaptic weight representing the risky cue in the patients increases rapidly to strengths that significantly exceed those of the controls in both models. In contrast, the D2 synaptic strength remains unchanged in the H2 model relative to the controls. At first glance, this seems counter intuitive given that H2 includes impaired D2 plasticity (increased LTD to LTP) however this lack of build-up of D2-Indirect pathway activity *is* pathological and reflects the blunted plasticity response to the negative prediction error. This can be understood when the D2 synaptic changes are compared between the H1 (Figure 6A) and H2 (Figure 6B) models. Under “physiological” D2 plasticity, the H1 model generates a substantial increase in D2-Indirect pathway activity which is proportionate to the increased risky choices and correspondingly increased negative prediction error signals that this risky choice behaviour produces. In contrast, in the presence of D2 synaptic plasticity, which is biased towards depotentiation under conditions of H2, there is no corresponding increase in indirect pathways weights and at a behavioural level, no time dependent devaluation of the risky cue. This difference between the two models suggests that for the combination of both reduced risk aversion and reduced choice devaluation observed in the DYT1 patients, cortico-striatal plasticity needs to be abnormal in opposing directions in both D1 and D2 Direct and Indirect BG pathways simultaneously.

**Figure 6:**
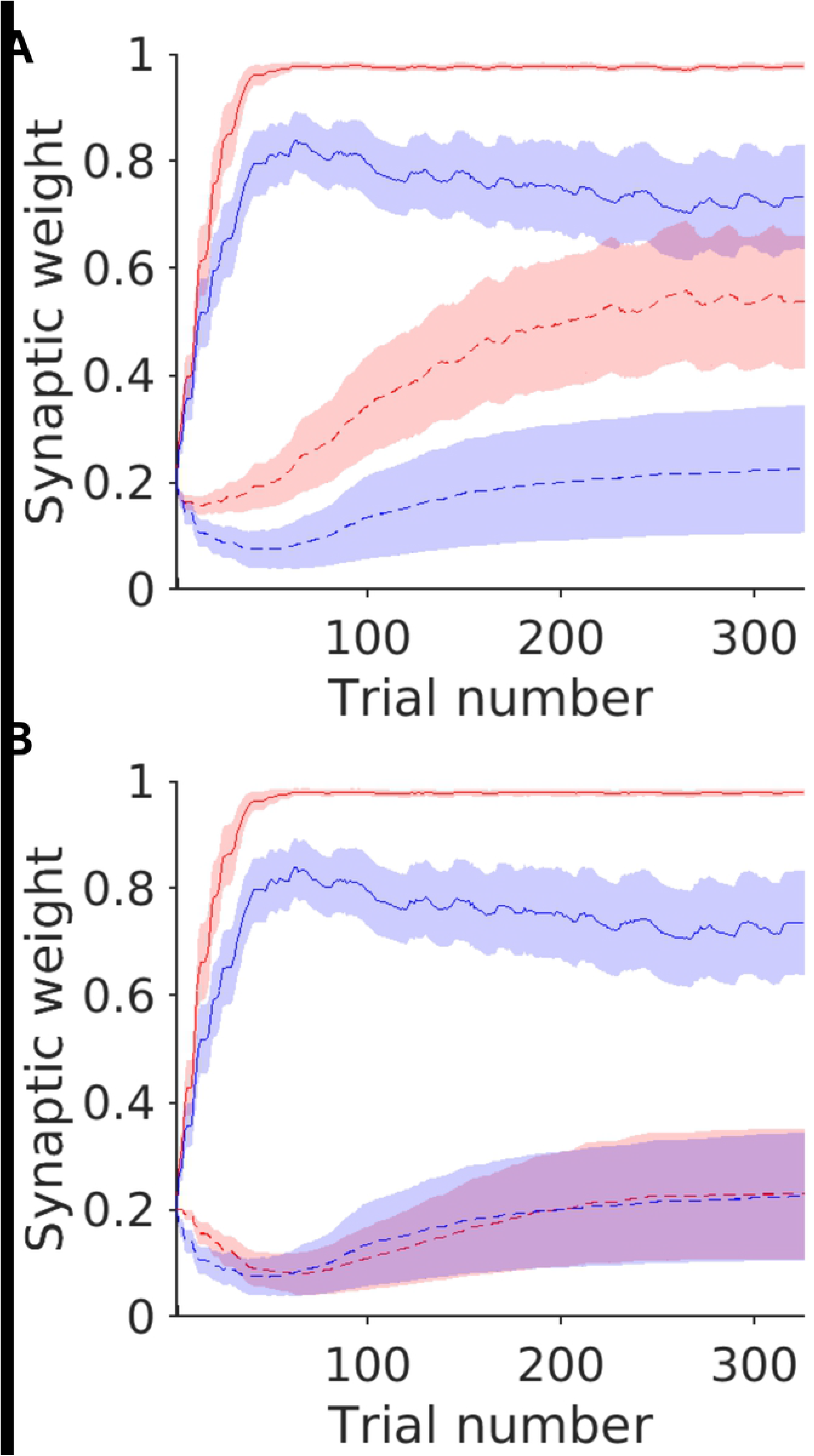
Simulated striatal synaptic weight changes during the task for the risky cue. Average simulated weights ± S.E.M (between simulations) for the CSP model under plasticity condictions of H1 (**A**) and H2 (**B**). Weights representing the risky cue are illustrated only. D2 weights, dashed lines (--) D1 weights, solid lines (-). Patients in red, controls in blue.

## Discussion

The purpose of this report was to highlight a disparity between the risk taking behaviour of patients with manifest signs of dystonia caused by the TOR1A mutation and increased cortico-striatal D2R-LTP observed *in vitro* from rodent models of human dystonia which do not manifest dystonic signs. Increased risk taking is a behaviour that is most likely explained by a combination of either impaired negative prediction error signalling (reduced D2R striato-pallidal LTP) or excess sensitivity of the positive prediction error signal (increased D1R striato-nigral LTP). Both of these lead to increased risk taking behaviour by making patients more likely to choose the “risky” cue following a “win”, or less sensitive to negative feedback following a “losing” choice. Using a simulation of cortico-striatal plasticity in the striatum, the risk taking performance levels of patients were least likely to be reproduced when the model incorporated increased D2R-LTP as reported from rodent experiments. The inferior performance of the model under these plasticity conditions was entirely attributable to the absence of D2-LTD, as increased D1-LTP was a feature of all three hypothetical plasticity abnormalities that were tested. When D2R plasticity was biased towards excessive LTD, the behaviour of the computational model was statistically indistinguishable from that observed experimentally by patients. This suggests the opposite pattern of D2R abnormalities to that reported in animal model *in vitro* experiments explains reinforcement learning abnormalities in patients observed by Arkadir et. al., (2016).

The reduced sensitivity to the negative prediction error signal in the DYT1 patients that contributes to increased risk taking is analogous to the reduced D2-LTD that underpins the reversal learning impairment seen in patients with cervical dystonia (Gilbertson et. al., 2019). These results support a common mechanism of increased D2-LTD causing abnormal reinforcement learning that is independent on the specifics of the task or clinical phenotype studied. As the density of D2R correlates with the sensitivity to negative decision outcomes (Frank *et al.*, 2009; Cox *et al.*, 2015) the loss of D2-LTP predicted by our model is also consistent with imaging studies demonstrating reduced D2R in both forms of dystonia (Naumann *et al.*, 1998; Carbon *et al.*, 2009).

Our approach was not intended to question the robustness of the method or interpretation of previous *in vitro* animal studies. It is important to emphasise that our opposing conclusions are not a *post-hoc* re-interpretation of the *in vitro* data on the basis of the outcome of our computational model simulations. Rather, the simulations provide evidence in support of our *a priori* hypothesis that loss of D2R-LTD plasticity is difficult to reconcile with our existing knowledge of the role of cortico-striatal plasticity in reinforcement learning. This discrepancy between the human and rodent D2R plasticity abnormalities have crucial implications for our understanding of DYT1 dystonia and the development of new therapies for patients with this condition. In the first instance, they support the idea that striatal neurochemistry is not in a state that is indifferent to dystonically manifest and non-manifest behavioural states. Notably, although animals with the TOR1A mutation have significant striatal neurochemical abnormalities, *they exhibit little to no phenotypic resemblance of a movement disorder*. It is conceivable therefore, that a reason for our results supporting an opposite pattern of D2R abnormal plasticity to that seen *in vitro*, reflects a difference between the manifesting dystonic and non-manifesting state. This explanation is supported by observations from previous studies. First, following the peripheral nerve injury necessary to induce dystonia-like posturing in TOR1A mutant rodents, these are accompanied by significant increases in striatal dopamine and decreases in D2R receptor expression (Ip *et al.*, 2016). This fundamental shift in dopaminergic neurochemistry has also been observed in recent post mortem studies comparing manifesting and non-manifesting carriers of the *TOR1A* mutant gene (Iacono *et al.*, 2019). Second, the study of Edwards et. al., (2006) emphasises the apparent paradox of how the same mutation can lead to an opposite physiological response depending on the clinically manifest state (Edwards *et al.*, 2006). Here TOR1A mutation carriers were tested using transcranial magnetic simulation protocols which induced LTD-like plasticity in healthy controls. These failed to induce any response in non-manifesting carriers but produced an exaggerated LTD-like response in the manifesting carriers.

Given this context it is unsurprising that our computational modelling of patient’s behaviour converges on a conclusion opposite to that reported from experiments using animal models of human DYT1 dystonia. Our results have crucial implications for the development of small molecular therapies based on translational studies in rodents (Downs *et al.*, 2019). We argue that since performance on reinforcement learning tasks correlates with the severity of the movement disorder in these patients, these tasks could be used to screen putative therapeutic agents based upon their ability to modify reward learning. This would be a cost effective intermediate step prior to formal clinical trial testing aimed to at the identification of novel agents.

## Acknowledgements

The authors thank Professor Mark Humphries for feedback on an earlier version of the manuscript.

## Author Contributions

TG analysed the data.TG, DA and DS wrote the paper. DA acquired performed the experimental paradigm and acquired the behavioural data.

